# A homozygous hypomorphic *BRCA2* variant causes primary ovarian insufficiency without cancer or Fanconi anemia traits

**DOI:** 10.1101/751644

**Authors:** Sandrine Caburet, Abdelkader Heddar, Elodie Dardillac, Helene Creux, Marie Lambert, Sébastien Messiaen, Sophie Tourpin, Gabriel Livera, Bernard S. Lopez, Micheline Misrahi

## Abstract

Primary Ovarian insufficiency (POI) affects 1% of women under forty. We studied a patient with a non-syndromic POI, from a consanguineous Turkish family. Exome sequencing identified a homozygous missense variant c.8524C>T/p.R2842C in *BRCA2*. BRCA2 is a major player in homologous recombination (HR). BRCA2 deficiency induces cancer predisposition and Fanconi Anemia (FA). Remarkably, neither the patient nor her family exhibit somatic pathologies. The patient’s somatic cells presented intermediate levels of chromosomal breaks, cell proliferation and radiation-induced RAD51 foci formation when compared to controls, the heterozygous mother’s and FA cells. R2842C-BRCA2 partially complemented BRCA2 depletion for double-strand break-induced HR. The residual HR function in patient’s cells could explain the absence of somatic pathology. BRCA2 is expressed in human fetal ovaries in pachytene stage oocytes, when meiotic HR occurs. This study has a major impact on the understanding of genome maintenance in somatic and meiotic cells and on the management of POI patients.

## INTRODUCTION

Primary ovarian insufficiency (POI) is a public health issue affecting ∼1% of women under 40 years, and is clinically heterogeneous with isolated or syndromic forms (Huhtaniemi et al., 2018). Most cases are idiopathic but an increasing number of genetic causes have been recently identified, especially mutations in genes involved in DNA repair and recombination (AlAsiri et al., 2015; Fouquet et al., 2017; Wood-Trageser et al., 2014).

DNA repair and recombination are essential for genome maintenance. The DNA damage response (DDR) coordinates a network of pathways insuring faithful transmission of genetic material. Consistently, defects in the DDR result in genome instability associated with developmental anomalies and cancer predisposition (Hoeijmakers, 2009). Homologous recombination (HR), an evolutionary conserved process essential to genome stability and cell viability, plays crucial roles in DNA double strand break (DSB) repair in somatic and meiotic cells.

In mammals, BRCA2 binds damaged DNA and loads the pivotal HR player RAD51, which then promotes DNA homology search. Therefore, cells defective in RAD51 or BRCA2 are thus defective in mitotic HR (Lambert and Lopez, 2000; Moynahan et al., 2001). Heterozygous *BRCA2* mutations increase susceptibility to breast and ovarian cancers, whereas severe bi-allelic defects in *RAD51* (*FANCR*) or *BRCA2 (FANCD1)* lead to Fanconi anemia (FA) syndrome (Tsui and Crismani, 2019). In particular, *FANCD1* syndrome associates developmental defects, genetic instability, bone marrow failure and cancer predisposition, with cancer developing in the first decade of life, and death before puberty (Meyer et al., 2014). The role of BRCA2 in RAD51 loading in mitotic HR makes it a strong candidate for an involvement in meiotic HR, but this remains to be formally established. Indeed, the severe phenotypes of bi-allelic inactivation of BRCA2 in humans and the early embryonic lethality resulting from germ-line inactivation of this essential gene in animal models hampered meiosis analysis and compromised the study of the putative functions of BRCA2 in gametogenesis (Ludwig et al., 1997; Sharan et al., 1997; Tsuzuki et al., 1996).

We describe here an adult patient carrying a homozygous missense mutation in *BRCA2* with isolated POI, but without cancer nor FA traits in the patient or her family. We demonstrate that the mutated R2842C-BRCA2 retains a lower but significant residual function when compared to wild-type (WT)-BRCA2. Consistently, the patient’s cells exhibit intermediate levels in chromosomal breaks, cell proliferation and ionizing radiation-induced RAD51 foci formation when compared to controls, a FANCD1 patient’s or the heterozygous mother’s cells. This residual HR in somatic cells could explain the absence of *in vivo* somatic pathologies. BRCA2 is a major cancer susceptibility gene and our finding will have a strong impact on the genetic counselling and management of patients with POI and their relatives.

## RESULTS

### Case report

The proposita was born to consanguineous Turkish parents (Figure 1). At 13 years, she had two vaginal bleedings followed by primo-secondary amenorrhea. She had normal pilosity, breast development and external genitalia. Several hormonal assays confirmed POI and pelvic ultrasonographical studies showed small ovaries with no or very few follicles (Table 1). Blood counts, liver and thyroid balances were normal. Thyroid auto-antibodies were undetectable. Bone densitometry at 30 years showed a marked osteopenia (T score = -2.3). She is presently 41 years old, displays normal blood assays and no other clinical sign. The karyotype is 46, XX and FMR1 premutation screening was negative. After two egg-donation procedures, she had two pregnancies with four healthy children.

**Table 1:** Clinical and biological studies of the proband and relatives. BMI: body mass index; SD: deviation compared to standards; R: Right; L: Left; FSH: Follicle-stimulating hormone; LH: Luteinizing hormone; E2: estradiol; AMH: anti-Müllerian hormone; InhB: inhibine B; T: testosterone; PRL: prolactin; TSH: thyroid-stimulating hormone.

| Case | Menstrual cycles | Age at evaluation (years) | BMI (height cm/weight kg) | Head circumference cm (SD) | (R/L) Ovarian Volume mm<sup>3</sup> | (R/L) Follicle number | FSH IU/l | LH IU/l | E2 nmol/l | AMH ng/ml | InhB ng/l | T nmol/l | PRL ng/ml | TSH mU/l |
| --- | --- | --- | --- | --- | --- | --- | --- | --- | --- | --- | --- | --- | --- | --- |
| Index | Primo-Secondary amenorrhea (13 years) | 30 | 23 (156/55) | 55.5 (0) | 21/17<br>0.9/0.5 | 2/0 | 85 | 16.7 | 0.04 |  |  | 0.5 | 17.5 | 1.45 |
|  |  | 32 |  |  | 19/17 | 0/0 | 113 | 36.5 | 0.08 |  |  | 0.6 | 29.5 | 1.34 |
|  |  | 39 | 24 (156/58) |  |  |  | 111 | 26.5 | 0.09 | 0.03 | < 8 | 1.43 | 4.6 | 1.66 |
| Mother | Secondary amenorrhea (33 years) | 63 | 38 (155/90) | 56 (0) | Normal ranges | Follicular phase | 2.9-12 | 1.5-8 | 0.06-0.54 | 2.2 - 6.8 | 10 - 320 | 0.4 - 2 | 0.7 - 25 | 0.3 - 4.2 |
|  |  |  |  |  |  | Ovulatory phase | 6.3-24 | 9.6-80 | 0.16-0.78 |  |  |  |  |  |
| Father |  | 70 | 17 (170/50) | 55 (0) |  | Luteal phase | 1.5-7 | 0.2-6.5 | 0.34-2.1 |  |  |  |  |  |
| Brother |  | 45 | 28 (171/80) | 57 (0) |  | Menopause | 17-95 | 8-33 | ≤0.2 |  |  |  |  |  |

**Figure 1:**
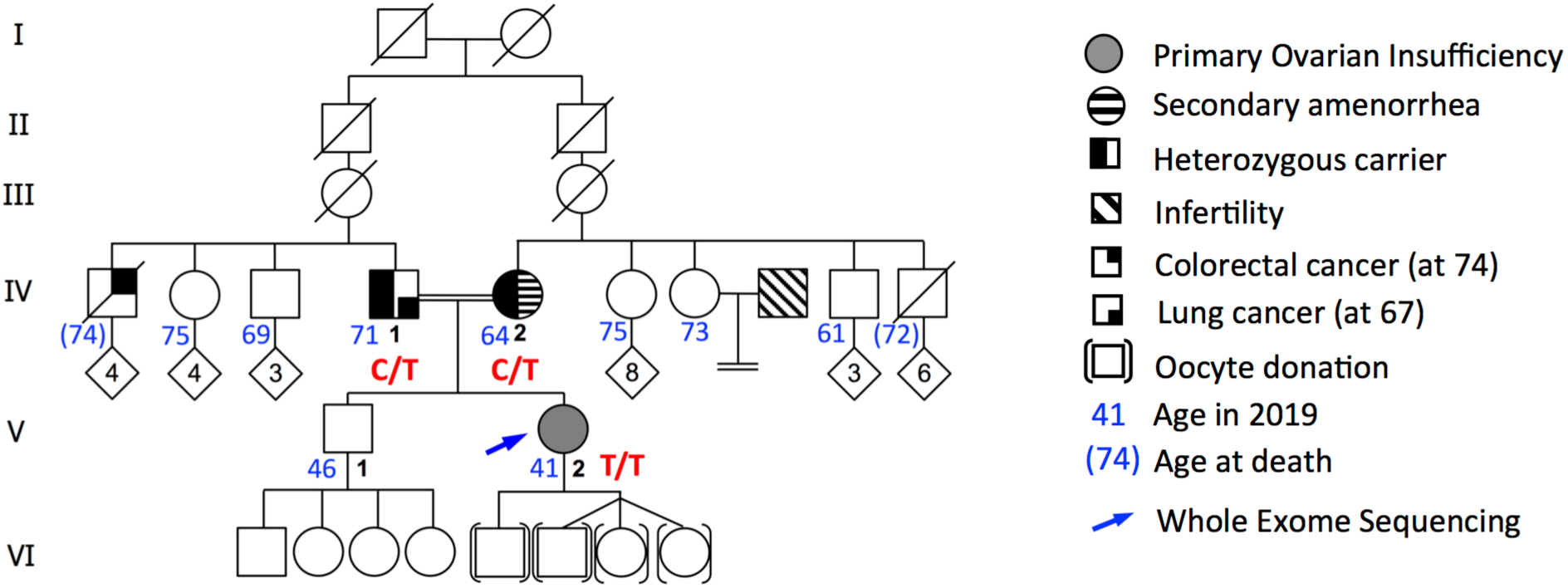
Pedigree of the Turkish family. Double lines indicate consanguineous union. The proband (blue arrow) was analysed by WES. The genotypes for the mutated codon of *BRCA2* are indicated in red.

The patient and family members have a normal stature, normal head circumferences with no abnormality in skin pigmentation, skeletal development or dysmorphia (Table 1). There was no familial history of infertility or other diseases. The mother married at the age of 15 and had two pregnancies at 18 (a boy) and 22 years (the proposita). This is in line with a delayed conception followed by secondary amenorrhea at the age of 33, not investigated in Turkey. She is obese with a BMI of 38. In order to rule out other genetic causes that could explain her subfertility, we performed a targeted NGS (see supplementary information). The brother had one healthy son (17 years) and 3 healthy daughters aged 15, 13 (both with normal puberty) and 8 years. The 71 years-old father, a heavy smoker, had a lung cancer at the age of 67 years, treated by radiotherapy and chemotherapy. A paternal uncle developed a colorectal cancer at the age of 74 years and died few months later. Six other paternal and maternal uncles and aunts are 61 to 75 years old and have no history of cancer or infertility.

### Whole-exome sequencing identified a homozygous missense variant in the DNA-binding domain of *BRCA2*

The patient was studied by whole-exome sequencing (WES). Familial consanguinity suggested an autosomal recessive inheritance pattern. The variants were therefore filtered on the basis of their homozygosity in the patient, their absence in unrelated fertile in-house controls and a minor allele frequency (MAF) below 0.01 in all available databases. Further filtering on available functional data for a possible role in fertility revealed the missense variant rs80359104, NM_000059.3: c.8524C>T (p.R2842C), located in exon 20 of *BRCA2* (Figure 2A, Figure 2 – figure supplement 1 and Figure 2–figure supplement 2). The variant is very rare and presents only at the heterozygous state in 3 out of 138342 individuals without known phenotype (MAF 1.10^-5^) in the non-cancer GnomAD subset. It is absent in the Greater Middle East Variome database dedicated to Middle Eastern populations. It is predicted to be pathogenic by 16 of 17 predictive softwares (Figure 2–figure supplement 3).

**Figure 2:**
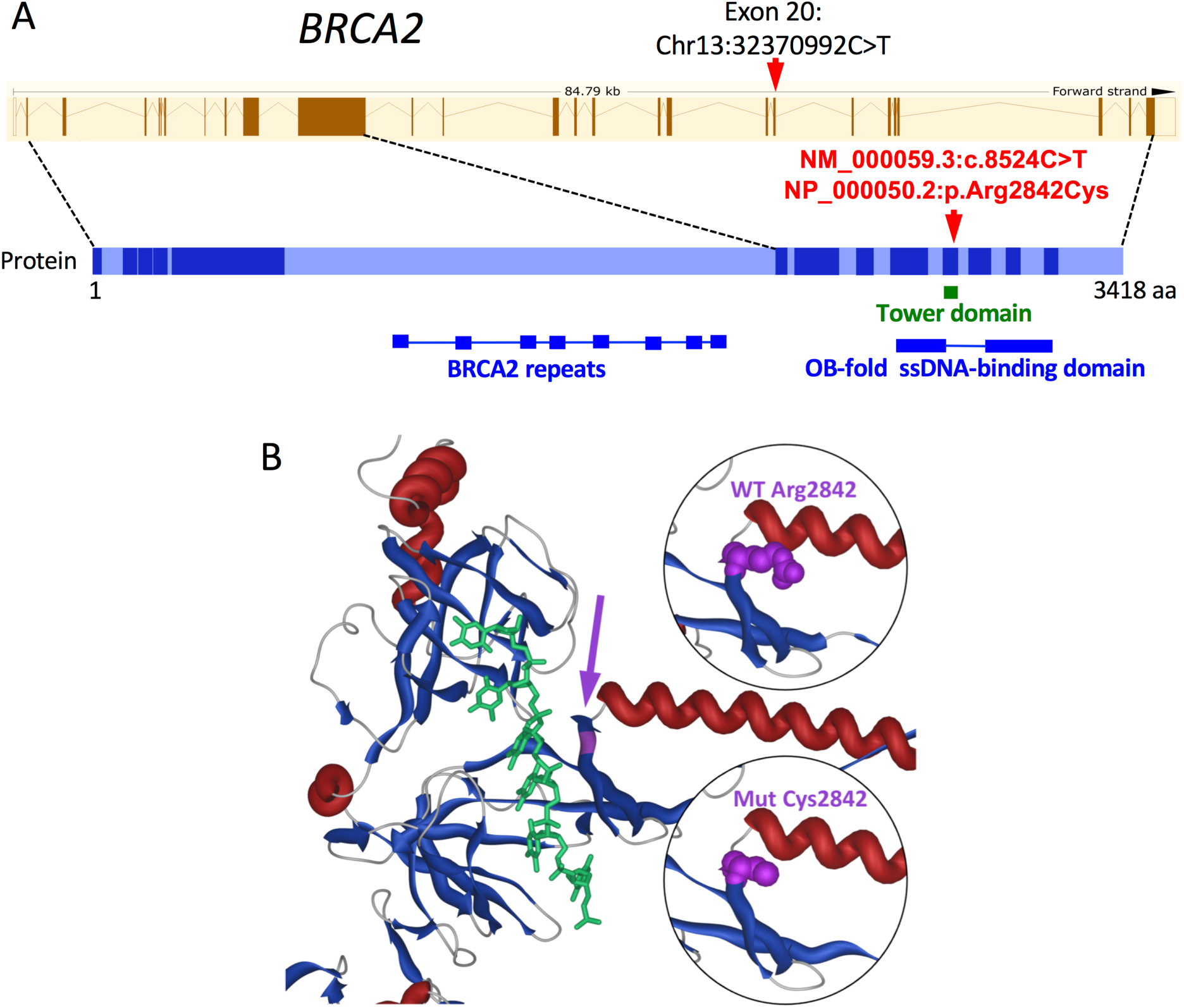
Mutation of *BRCA2* in a POI patient without FA trait. **A.** Position of the variant in *BRCA2* gene and protein. The structure of the normal protein for the longest isoform of 3418 residues is shown below the genomic structure with the coding exons as coloured bars (Ensembl, reference transcript ENST00000544455.5). The mutation (red arrow) lies at the very end of exon 20, which encodes 48 aminoacids (aa) encompassing the Tower domain at the center of the OB-fold ssDNA-binding domain (oligonucleotide/oligosaccharide-Binding single-strand DNA-binding domain). **B.** Partial view of the 3D model of the BRCA2 C-terminal domain (alpha-helices in red, beta-sheets in blue). The mutated position (purple) is located near the ssDNA (green), at the base of the Tower domain, that forms a stem of two long alpha-helices and a helix-turn-helix motif, similar to the DNA-binding domains of recombinases and homeodomain transcription factors. Inserts: difference between the occupancy of the lateral chain of wild-type (WT, top) and mutated (bottom) residue at this position. Figure 2–figure supplement 1: WES metrics for the POI patient Figure 2–figure supplement 2: Filtering of the variants identified in the POI patient Figure 2–figure supplement 3: Pathogenicity predictions for the R2842C variant in BRCA2 Figure 2–figure supplement 4: Conservation of the mutated Arg 2842 across species.

The variant changes a strictly-conserved aminoacid (aa) at the base of the Tower part of the BRCA2 DNA-binding domain, in close proximity to the groove that binds single-stranded DNA (ssDNA) (Figure 2B and Figure 2–figure supplement 4). This C-terminal domain is essential for appropriate binding of BRCA2 to ssDNA (Yang et al., 2002).

BRCA2 loads recombinases on ssDNA: RAD51 in mitotic cells, RAD51 and the meiotic DMC1 in germ cells, indicating a crucial role for BRCA2 in mitotic and meiotic HR (Martinez et al., 2016). Cells defective in RAD51 or BRCA2 are defective in mitotic HR (Lambert and Lopez, 2000; Moynahan et al., 2001) and in mouse, germ cells-specific *Brca2* deletions lead to meiotic impairment and infertility (Miao et al., 2019; Sharan et al., 2004). Therefore, we considered *R2842C*-*BRCA2* as a very likely causal variant for isolated POI in our patient, although she does not present FA traits. Hence, we investigated the expression of *BRCA2* in human oocytes and the functional impact of the variant on HR in human cells.

### *BRCA2* is expressed during meiotic prophase I in human fetal ovaries

In order to support a possible role for BRCA2 in female meiotic HR that would explain this patient’s infertility, we verified its expression and localisation in human fetal ovaries. Indeed, *Brca2* mRNA expression was reported in murine oocytes (Sharan et al., 2004) and BRCA2 was described to form recombination nodule-like foci along chromosome axes in human spermatocytes (Chen et al., 1998), but its expression during human female meiosis remained undocumented. Using qRT-PCR on RNA libraries prepared from human ovarian samples at various fetal stages, we detected a predominant expression of *BRCA2* mRNA after 11 weeks post-fertilization, when oocytes enter and progress through meiotic prophase I (Figure 3A).

**Figure 3:**
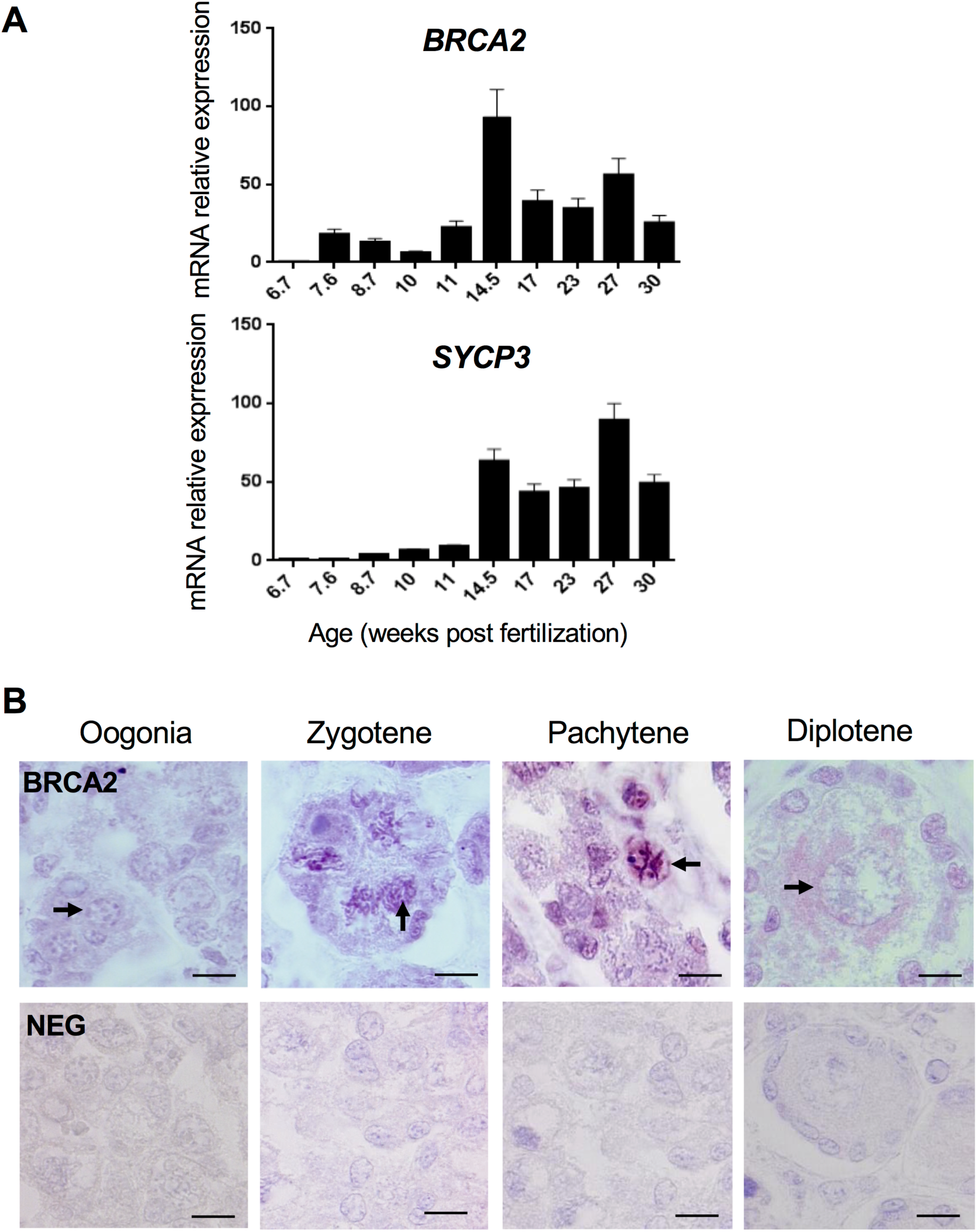
BRCA2 expression in human fetal ovaries. **A.** *BRCA2* mRNA was quantified by RT-qPCR from total RNA of pooled human fetal ovaries from various developmental stages (above). Beta-actin was used as a reference and expression is provided as percentage of the maximum. Each RNA library was analysed in triplicate and the bar indicates the mean. Quantification of *SYCP3* mRNA is shown at the same stages for comparison (below). **B.** BRCA2 immunostaining (purple) in the cortex of a 24 weeks post fertilization human ovary. Meiotic chromosomes are stained in zygotene and pachytene stage oocytes. No staining is observed when immunohistochemistry is performed in the absence of primary antibody (NEG). Arrowheads point to germ cells at the indicated stage. Scale bar: 10 µm.

Immunostaining of human fetal ovarian sections showed that BRCA2 protein was detected mostly in pachytene stage oocytes (Figure 3B). BRCA2 staining appeared as thick threads, likely corresponding to meiotic chromosomes. These results show that BRCA2 is indeed present on chromosomes in fetal human oocytes when meiotic DSB repair occurs.

### *R2842C-BRCA2* displays a reduced DSB-induced HR efficiency

Although referenced in the COSMIC database of variants in cancer (COSM23938), *R2842C-BRCA2* is considered as a variant of unknown significance for breast cancer predisposition (VUS, IARC class 3). Attempts to classify BRCA2 VUS in a hamster lung fibroblast cell line showed that this variant displayed a little decrease in HR efficiency, at the limit for inferring pathogenicity (Guidugli et al., 2013). Therefore, the significance of this variant and its classification as a causal mutation for human pathology remained unclear. A previous attempt to classify BRCA2 VUS in a hamster lung fibroblast cell line showed that this variant displayed a little decrease in HR efficiency, at the limit for inferring pathogenicity (Guidugli et al., 2013). Since human and rodent cells differ in their regulation of DSB repair, we analysed the specific impact of R2842C-BRCA2 on HR in human cells. We used the RG37 cell line (Dumay et al., 2006), a human SV40 immortalized fibroblast line bearing the DR-GFP substrate (Pierce et al., 1999) that monitors gene conversion induced by targeted cleavage by the I-SceI meganuclease (Figure 4A). Both the expression of WT-BRCA2 and R2842C-BRCA2 stimulated the efficiency of DSB-induced HR (Figure 4B). We then silenced the endogenous *BRCA2* using a specific siRNA targeting its 3’UTR sequence, and complemented these cells with either the WT or mutated BRCA2 (Figure 4C). As expected, silencing endogenous *BRCA2* decreased HR efficiency, and WT-BRCA2 fully complemented HR efficiency. R2842C-BRCA2, expressed at similar levels than WT-BRCA2, only partially complemented HR efficiency, to 67 ± 6% compared to WT-BRCA2 (Figure 4B).

**Figure 4:**
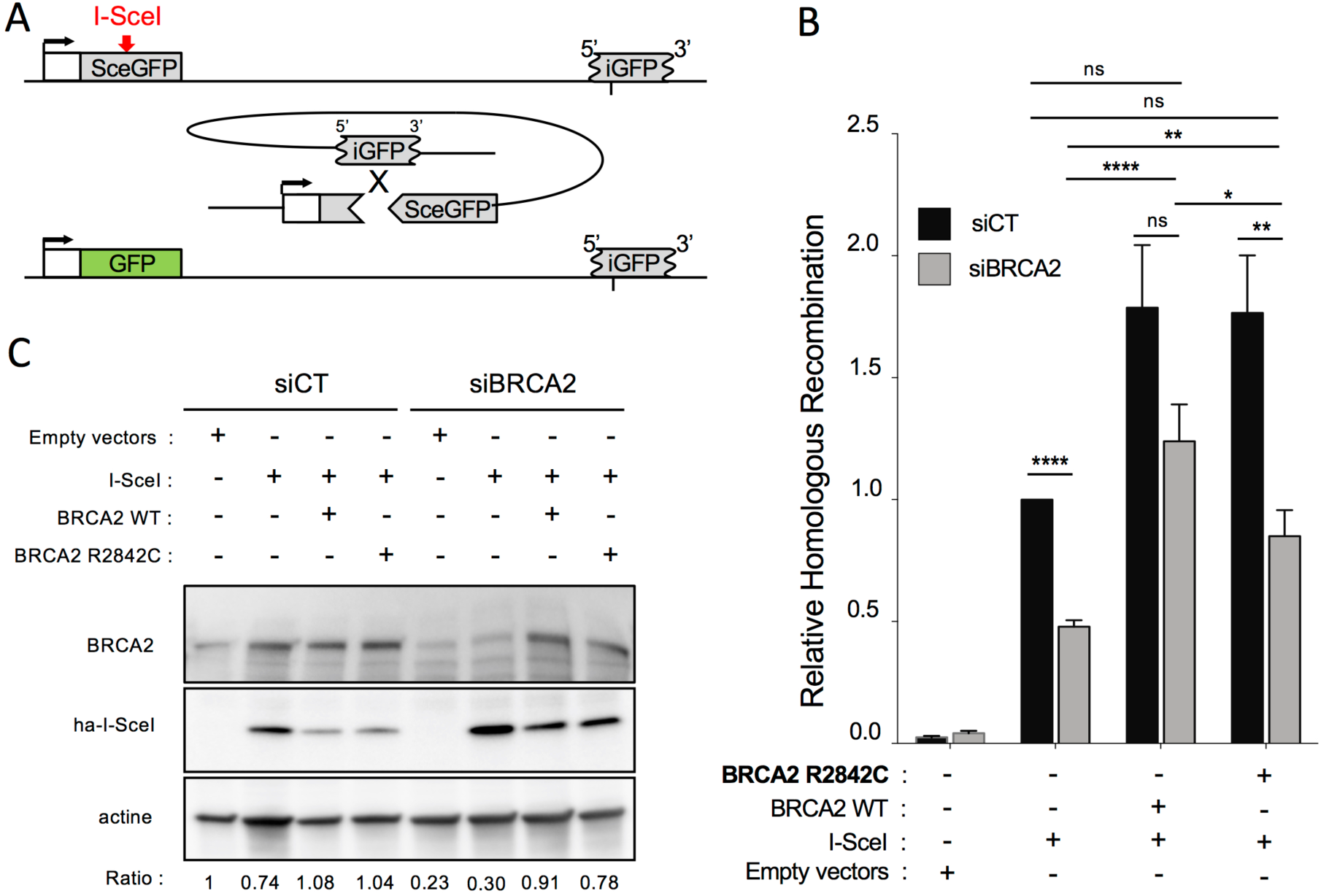
Impact of the *R2842C-BRCA2* mutation on homologous recombination induced by targeted DSB. **A.** Schematic representation of the DR-GFP substrate for the study of homologous recombination. Two inactive GFP (iGFP and SceGFP) genes are organised into direct repeats. The I-SceI meganuclease generates a targeted DSB cleavage into the substrate (red). HR between the two GFP genes generates a functional GFP. The DR-GFP substrate is stably integrated in the SV40-transformed fibroblasts RG37 cell line, and the relative HR efficiency is quantified as the fraction of GFP-positive cells (i.e with a repaired GFP gene after targeted cleavage), as scored by FACS. **B.** HR efficiency, measured by the fraction of GFP+ cells, in cells expressing the WT or mutated BRCA2 protein (normalised to I-SceI transfected cells), and transfected either with a control siRNA (siCT) or a siRNA targeting the 3’UTR of endogenous *BRCA2* mRNA (siBRCA2). The values are normalised to the control and represent the average ± SEM (p-values from Mann-Whitney test) for at least 3 independent experiments. **C.** Expression of endogenous BRCA2 and exogenous WT-BRCA2 and R2842C-BRCA2. Twenty micrograms of total proteins extracted from a wild type or R2842C mutant BRCA2-expressing cell line were electroblotted in the presence of endogenous BRCA2 (siCT) or after specific silencing (siBRCA2). For each condition, the expression of I-SceI and BRCA2 and the efficiency of silencing were measured. We used β-actin as a loading control. Below: relative quantification of BRCA2 versus actin by quantification of bands intensity with ImageJ.

These data show that the *R2842C-BRCA2* mutation affects HR efficiency in human cells, but only partially. This significant residual activity could account for the absence of somatic pathology in the patient.

### Increased chromosomal instability in the patient’s cells

Then we studied mitomycin C (MMC)-induced chromosomal breaks in lymphoblastoid cells derived from the patient, her mother, two fertile control women and a FANCD1 patient. In the absence of MMC, few spontaneous breaks were observed in cells from the proposita and the FANCD1 patient (Figure 5A and 5B). Upon exposure to 300 nM MMC, all FANCD1 cells presented breaks, as expected, while the patient’s cells exhibited a slight increase of chromosomal breaks, compared to the heterozygous mother’s and the WT control cells. Furthermore, at high MMC dose (1000 nM), while breaks in FANCD1 cells were too numerous to be quantified, the patient’s cells presented only a modest increase of breaks (Figure 5B). These data show that the patient’s cells display levels of chromosomal breaks intermediate between those of WT and FANCD1 cells.

**Figure 5:**
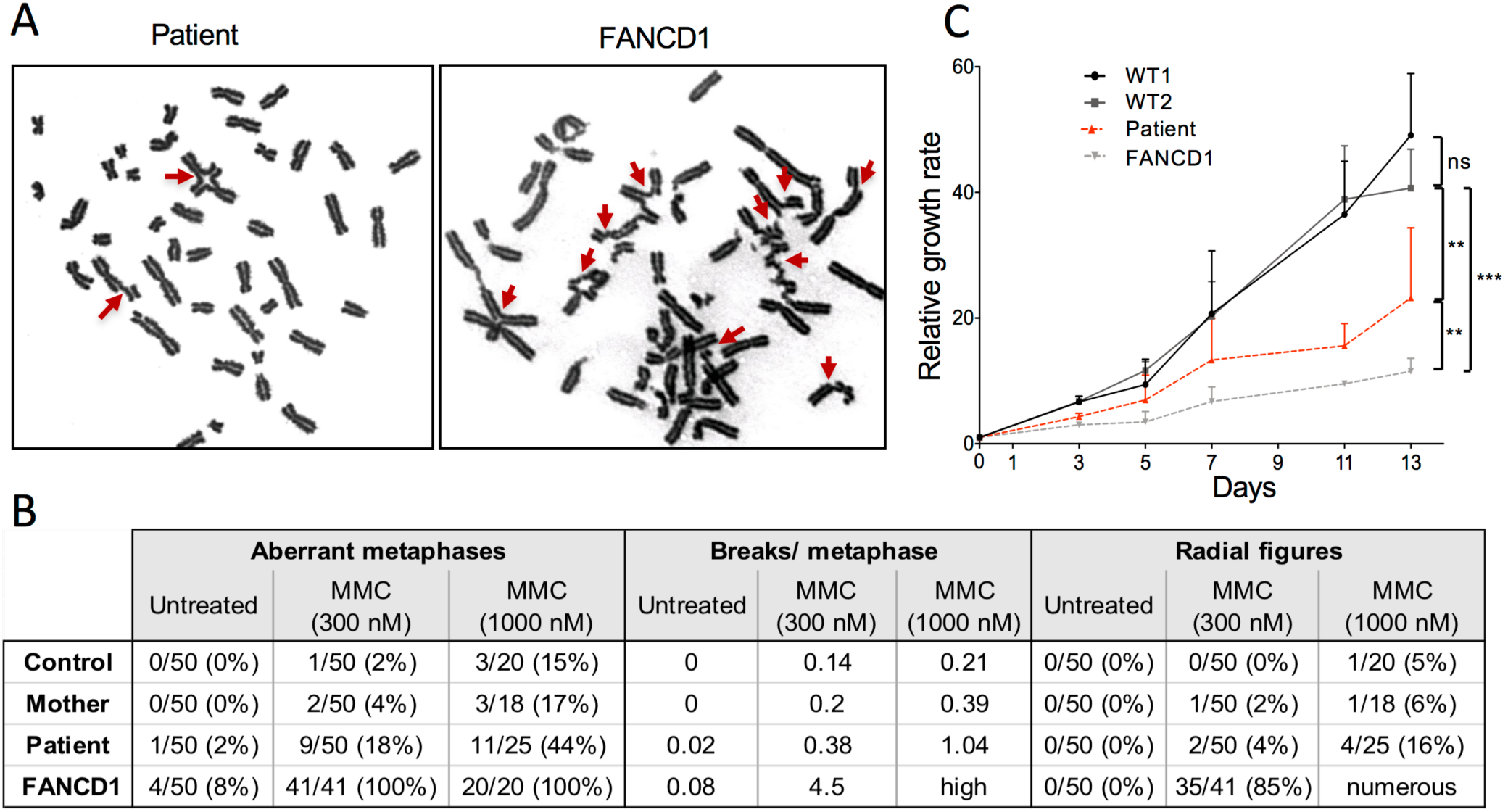
Increased chromosomal instability and reduced proliferation rate in the patient’s cells. **A.** Chromosomal breaks analysis of lymphoblastoid cell lines. Metaphases in the patient’s and FANCD1 cells in the presence of 300 nM Mitomycin C (MMC). Chromosomal breaks and radial figures are shown (arrows). **B.** Quantification of chromosomal breaks in lymphoblastoid cell lines derived from the patient, the mother, a *FANCD1* patient and a WT control, in the absence or in the presence of MMC. **C.** Reduced proliferation rate of the patient’s primary fibroblasts. Cells from the POI patient, two WT controls (WT1 and WT2) and a FANCD1 patient were cultured into 6-wells plates and counted every 2-3 days during thirteen days. The value corresponds to the mean + SEM of at least 3 independent experiments. The statistical significance was calculated from rope comparison of linear regression of growth curves.

### Reduced proliferation rate of the patient’s primary fibroblasts

Then we compared the proliferation rate of primary fibroblasts from the POI patient, WT controls and a FANCD1 patient. As expected, the proliferation of FANCD1 cells was markedly affected when compared to that of the WT cells (Figure 5C). Remarkably, the patient’s fibroblasts exhibited a moderately reduced proliferation rate, intermediate between the WT and the FANCD1 cells.

### Altered radiation-induced RAD51 foci formation in the patient’s fibroblasts

The main role for BRCA2 is the loading of the pivotal RAD51 recombinase on damaged DNA, a crucial step for triggering HR. Therefore, we monitored the radiation-induced assembly of RAD51 foci, which are considered sites of HR initiation events. As expected, FANCD1 cells failed to assemble RAD51 foci (Figure 6A and 6B). The patient’s cells showed an intermediate phenotype: indeed, at 6 Gy, they assemble foci with kinetics comparable to WT cells, but the level of the plateau was about two-fold lower than in WT cells (Figure 6B). A dose-response analysis confirmed the complete deficiency in RAD51 foci assembly in FANCD1 cells, at all irradiation doses (Figure 6C and 6D). In the patient’s cells, the number of RAD51 foci increased up to 2 Gy similarly to WT cells, but did not further increased at higher doses (Figure 6C, Figure 6D left panel). Consequently, the patient’s cells showed lower levels of RAD51 foci at high doses (>2 Gy) when compared to WT cells (Figure 6C, Figure 6D right panel). Together, these data show a dose-dependent sensitivity in the patient’s cells, able to process low levels of DNA damage, but failing to assemble RAD51 foci when faced with high levels of damage.

**Figure 6:**
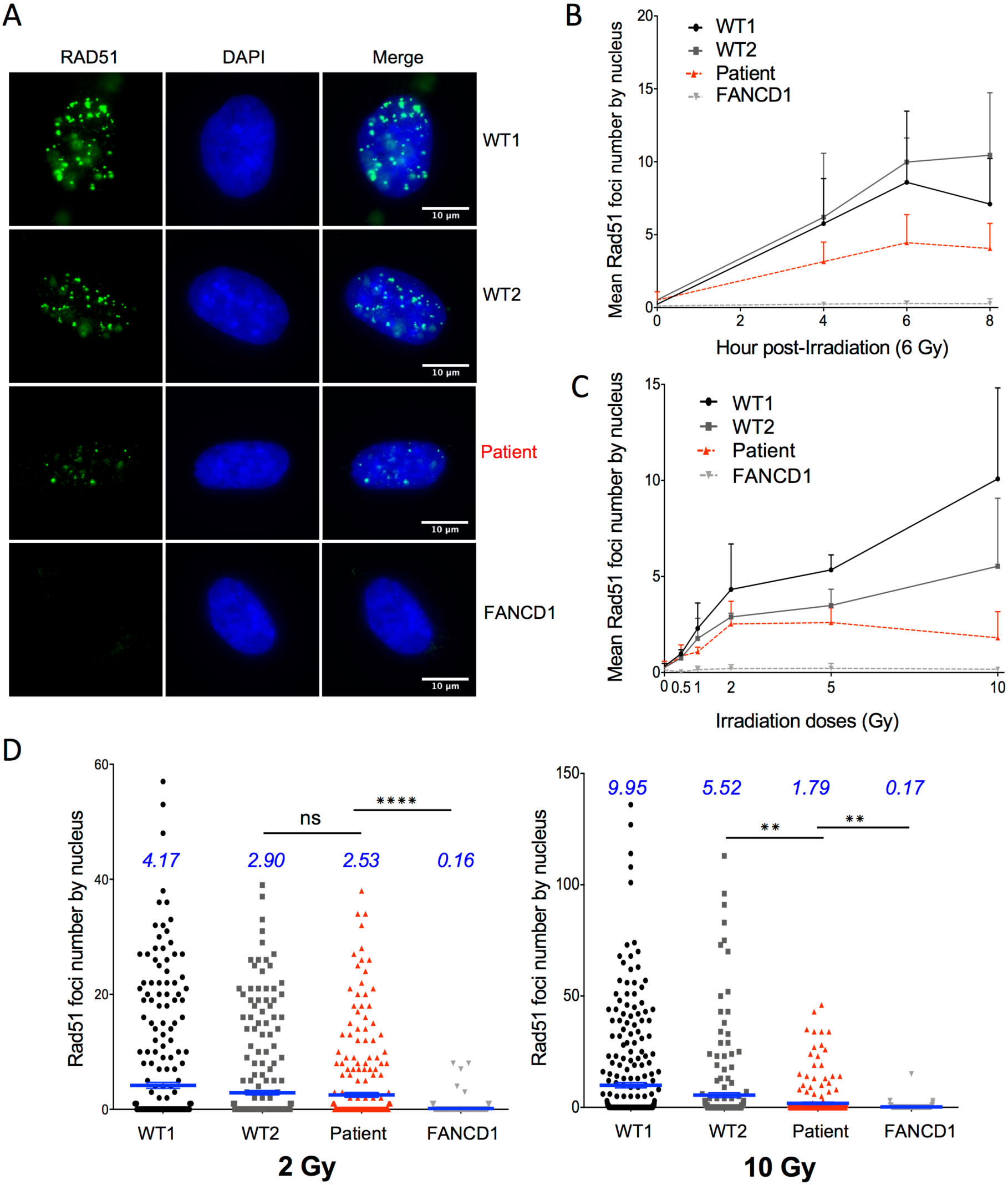
Dose-sensitive alteration of RAD51 foci formation in the patient’s cells. **A.** RAD51 nuclear foci assembly in irradiated primary fibroblasts from the patient, two WT controls and a FANCD1 patient (6 Gy, 6 hours). Fixed and permeabilised cells were probed with an anti-RAD51 antibody. **B.** Kinetics of RAD51 foci assembly (6 Gy) in irradiated primary fibroblasts. **C, D.** Dose response of RAD51 foci assembly (6h post-irradiation) in irradiated primary fibroblasts. **C.** Complete dose response of RAD51 foci per nucleus; **D.** RAD51 foci at low (2 Gy, left panel) and high (10 Gy, right panel) levels of irradiation respectively (medians are in blue (n=3)). In B and C, the values correspond to the mean + SEM (n=3); p-values (stars) were obtained with Mann-Whitney test. Figure 6-figure supplement 1: Compilation of WES data for all genes involved in the FA pathway

Bi-allelic mutations in two distinct genes of the FA pathway can be found in a single individual (Singh et al., 2009). In order to rule out that the cellular phenotypes observed in the patient’s cells could be due to mild bi-allelic mutations in FA genes other than *BRCA2*, we have analysed the variants found by WES in the 26 other FA genes. The good coverage of each gene, the presence of heterozygous variants (both in coding regions and in UTRS or intronic regions) and unbiased allelic ratios exclude the possibility of having missed a pathogenic variant in another FA gene (Figure 6-figure supplement 1). This analysis supports the fact that the cellular phenotypes observed in the patient’s cells are not due to mutations in FA genes other than *BRCA2*. In particular, no variants in *RAD51* could explain the defects in foci assembly detected in the patient’s cells.

## Discussion

We report here for the first time a *BRCA2* homozygous hypomorphic mutation in a patient with POI, and, remarkably, without cancer nor Fanconi anemia traits in the patient and her family. The mutation is located in the ssDNA-binding domain of BRCA2. We performed a thorough functional study and showed that the R2842C-BRCA2 mutant displays a reduced residual HR activity. Consistently, the patient’s cells exhibit a reduced rate of proliferation, a slight increase in MMC-induced chromosomal breaks and a dose-sensitive alteration of radiation-induced assembly of RAD51 foci. However, despite these moderate alterations in somatic cells, the patient did not develop somatic pathology, unlike FANCD1 patients.

Meiotic recombination is a complex and highly regulated process occurring in meiotic prophase I and involving specific meiotic genes such as *DMC1* and *MEIOB* (Crickard et al., 2018; Souquet et al., 2013; Yoshida et al., 1998). While the role and clinical impact of homozygous and heterozygous *BRCA2* variants in mitotic cells is widely studied, the impact of such defects in human germ cells remains less understood because almost all FANCD1 patients die before puberty. However, it was very recently shown in mouse models that oocyte-specific *Brca2* defects lead to meiotic impairment, germ cells depletion and infertility (Miao et al., 2019; Tsui and Crismani, 2019). Here, we show that BRCA2 mRNA and protein are expressed in human fetal ovaries in pachytene stage oocytes, when meiotic HR occurs. Taken together, our data strongly support an actual role of BRCA2 in meiotic HR, and therefore a likely impact of R2842C-BRCA2 reduced activity on this process in our POI patient.

The main role of BRCA2 is the loading of the pivotal recombinase RAD51 on damaged DNA, to allow for repairing DSBs by HR. Using a HR reporter assay and quantification of RAD51 foci in the patient’s irradiated somatic cells, we show here that the R2842C-BRCA2 mutant exhibits a reduced DSB-induced HR efficiency, but that RAD51 foci assembly was not affected at low irradiation doses. In addition, the patient’s cells displayed a growth rate and levels of chromosomal breaks intermediate between FANCD1 cells and WT cells. This shows that the patient’s somatic cells are slightly altered when compared to WT cells, but at a level insufficient to cause somatic pathological consequences. Since R2842C-BRCA2 cells efficiently load RAD51 on low numbers of DNA damage, the residual HR activity of the R2842C-BRCA2 mutant could account for the absence of somatic disorders.

Meiotic recombination in germ cells is initiated by hundreds of DSBs, and crossing-overs (involving HR) are required to produce functional gametes (Zhang et al., 2019). By contrast, somatic DSBs are introduced by accident and such cells are rarely spontaneously confronted to such high levels of simultaneous DSBs. In our functional test, RAD51 foci assembly by the mutated BRCA2 was significantly decreased at higher doses (>5 Gy) that generate a high number of DSBs (with an estimation of 30 to 40 DSB/Gy/mammalian genome (Ruiz de Almodóvar et al., 1994)). Therefore, the *R2842C-BRCA2* mutation is expected to affect the processing of such a high number of simultaneous meiotic DSBs, explaining the infertility observed in our patient.

Heterozygous BRCA2 mutations increase susceptibility to familial breast and ovarian cancer (Walsh et al., 2011). Such increased susceptibility has not been observed in our patient’s family. The efficient loading of RAD51 on low number of DNA damages in the patient’s somatic cells could explain the absence of somatic pathologies. Although we cannot rule out the possibility that the patient may develop cancer on the long-term, after triggering by environmental causes, the fact that neither the patient nor her relatives have yet developed other pathologies implies that the *BRCA2* mutation is hypomorphic and retains a residual HR activity, as supported by our functional studies.

Recently, two young sisters presenting a syndromic XX ovarian dysgenesis were reported to carry compound heterozygous C-terminal *BRCA2* truncations. However, in addition to POI, these patients and their family fulfilled the diagnostic criteria of FA: microcephaly, café-au-lait spots, childhood leukemia, and a characteristic severe cellular response to mitomycin in chromosomal breakage tests (over 100 breakages per cells at 150 and 300 nM MMC, 50 times the number observed in control lymphocytes), the hallmark of FA (Auerbach, 2009; Weinberg-Shukron et al., 2018). Therefore, these cases are not similar to our patient that presents only an isolated POI.

In conclusion, we describe and functionally characterize here for the first time a homozygous hypomorphic variant of BRCA2, in a patient with isolated POI without somatic pathology in the patient and her family. The recent implication of DNA repair genes in POI establishes a genetic link between infertility and cancer. As *BRCA2* is a major susceptibility gene for breast and ovarian cancer, this represents a major ethical issue for the care of these patients. It should change the genetic counselling and pre-test information for patients with isolated POI and their families. Indeed, such counselling should be addressed while keeping in mind a possible defect in major DNA repair genes such as *BRCA2*. More generally, this study has also a wide impact for the understanding of the processes controlling genome plasticity and the consequences of their defects, in somatic and germ cells.

## MATERIAL AND METHODS

### Ethics statement

The study was approved by all the institutions involved and by the agence de Biomedecine (reference number PFS12-002). Written informed consent was received from participants prior to inclusion in the study.

### Whole Exome Sequencing and bioinformatics analysis

WES, reads quality check and mapping was performed by Beckman Coulter Genomics (Danvers, USA). Exon capture was performed using the hsV5UTR kit target enrichment kit. Mapping was performed on the GRCh37.p13 reference genome using the Burrows-Wheeler Alignment tool (BWA) version 0.6.1-r104 with default parameters, and samtools ‘Rmdup’ was used to remove duplicates. Prior to variant calling, reads were re-aligned around known or suspected indels by the GATK Realigner commands. The samtools version 2.0 ‘mpileup’ command and the bcftools multi-allelic calling model were used for variant calling. Variants were annotated by SnpEff, VEP (Variant Effect Predictor) and dbNSFP 3.5a. Minor Allele Frequencies were manually verified using ExAC (http://exac.broadinstitute.org/), Gnomad (https://gnomad.broadinstitute.org/) and Kaviar (http://db.systemsbiology.net/kaviar/) databases.

### Sanger Sequencing Analysis

To confirm the presence and segregation of the variant, direct genomic Sanger DNA sequencing of *BRCA2* was performed in the patient and both parents using specific *BRCA2* primers: 5’-GACTACCCTCTCATAGCTCCAG-3’ and 5’-GGAAGAAGCAGGGAACACTC-3’

### Protein modelisation

The 1mje and 1miu structures of BRCA2 were retrieved from PDB databank 1. The 1mje structure was opened in iMol (Piotr Rotkiewicz, “iMol Molecular Visualization Program,” (2007) http://www.pirx.com/iMol) for displaying the proximity of the mutation to the DNA-binding groove in the C-terminal domain of the BRCA2 protein. The human WT and mutated sequence were threaded onto the 1miu structure using RaptorX 2, and the resulting pdb structures were rendered in iMol for displaying the occupancy of lateral chains.

### Collection of human samples

GM3348 (WT1) and GM3652 (WT2) are wild-type primary human fibroblasts (Coriell institute, Camden, USA). EGF 208_F, noted as FANCD1 cells, are primary fibroblast from a FANCD1 patient biopsy (generous gift from Dr. Jean Soulier, Hopital St Louis, Paris). Primary fibroblasts were derived from a skin biopsy of the patient. EBV-immortalized lymphoblastoid cell lines derived from the patient, both parents and two healthy women as control were established at the Banque de cellules, Genopole (Evry, France) using a standard protocol.

### Collection of human fetal gonads

Human fetal ovaries were obtained and studied as described (Frydman et al., 2017). Fetal ovaries were harvested from material obtained following legally induced abortions or therapeutic terminations of pregnancies at the Department of Obstetrics and Gynecology at the Antoine Béclère Hospital, Clamart (France). All women provided an informed consent and this study was approved by the Biomedicine Agency (reference number PFS12-002). Fetal age was calculated by measuring the length of limbs and feet according to a developed mathematical model (Evtouchenko et al., 1996). After collection, fetal gonads were stored in RLT RNA lysis buffer (Qiagen, Courtaboeuf, France) for gene expression profiling or fixed for histology and immunostaining. Fetal ovaries from the therapeutic terminations of pregnancies (second and third trimester of pregnancy) had to display normal histological features before being included in the study.

### Detection of BRCA2 in human fetal ovaries

Immunohistochemistry was studied as previously described (Poulain et al., 2015). Fetal human ovaries were fixed overnight in 10% neutral formalin (Carlo Erba Reagents, Val de Reuil, Frane) before being dehydrated, embedded in paraffin wax and cut into 5µm sections. After dewaxing and rehydration, antigen retrieval was performed in HIER citrate buffer pH 6 (Zytomed, Diagomics, Blagnac, France) in an autoclave (Retriever 2100, Proteogenix, Mundolsheim, France). Sections were then bathed in distilled water and incubated for 15 min in 3% H2O2 at room temperature. After 30 min in 2.5% normal Horse serum (Vector laboratories, Eurobio, Les Ulis, France), primary antibody diluted in PBS was incubated for 2h at 37°C. The primary antibody used in this study was rabbit polyclonal to human BRCA2 (1:200, Abcam, Paris, France). The primary antibody was revealed using the secondary antibody anti-rabbit IgG (IMPRESS kit, Vector Laboratories, Eurobio). Peroxidase activity was visualized using VIP (Vector laboratories, Eurobio) as a substrate. Sections were counterstained with hematoxylin.

### Real-time quantitative PCR

In order to measure the expression of multiple genes during human gonadal development, total RNA from fetal ovaries was extracted using the RNeasy Mini Kit (Qiagen Courtaboeuf, France), followed by a reverse transcription and whole transcriptome amplification (Quantitect Whole Transcriptome cDNA Amplification, Qiagen, Courtaboeuf, France). Seventeen ovaries were included for gene expression profiling as previously described (Poulain et al., 2014). Each RNA sample was analysed in triplicate. The 7900HT Fast Real-Time PCR System (Applied Biosystems, Foster City, CA) and SYBR-green labelling were used for quantitative RT-PCR. The comparative ΔΔcycle threshold method was used to determine the relative quantities of mRNA using *ACTB* (ß-actin) mRNA as reference gene for normalization. The sequences of oligonucleotides used with SYBR-green detection were designed with Primer Express Software:

ACTB: 5’-TGACCCAGATCATGTTTGAGA-3’; 3’-TACGGCCAGAGGCGTACAGG-5’

BRCA2: 5’-AGACTGTACTTCAGGGCCGTACA-3’; 3’-GCTGAGACAGGTGTGGAAAC-5’.

SYCP3: 5’-TGCGGTGTGTTTCAGTCAGG-3’, 3’-TTTTTCCGGAGGACACCATATT-5’

### Chromosome breakage studies

Chromosome breakage studies were performed in EBV-immortalized lymphoblastoid cell lines derived from the patient, her mother, a FANCD1 patient and a healthy woman as control. They were studied at the Gustave Roussy Institute (Villejuif, France), following a standard in-house protocol. EBV-immortalized cells were cultured under standard conditions for karyotyping. DNA damage was induced using Mitomycin C (MMC, Sigma) added for 48h. For each sample, three conditions were tested: without MMC to analyze spontaneous damages, and with 300 nM and 1000 nM MMC. Chromosome breakages were scored by an experimented cytogeneticist on at least 20 metaphases.

### Cell proliferation assay

Primary fibroblasts from the POI patient, two healthy WT controls (GM3348 and GM3562) and a FANCD1 patient were seeded into 6-well cell culture and grown in MEM (Gibco, Life Technologies) supplemented with 20% fetal calf serum (FCS; Lonza Group, Ltd.) and were incubated at 37°C with 5% CO_2_. For thirteen days, cells were dissociated from wells with trypsin and counted every 2-3 days using a Z1 Particle Counter (Beckman Coulter).

### Cell transfection and HR efficiency test

The HR efficiency was assessed in RG37 cell line, derived from SV40-transformed GM639 human fibroblasts in which we stably integrated the pDR-GFP gene conversion reporter (Dumay et al., 2006; Pierce et al., 1999). RG37 were cultured in DMEM supplemented with 10% fetal calf serum (FCS) and 2 mM glutamine and were incubated at 37°C with 5% CO_2._ For HR efficiency test, The I-SceI meganuclease was expressed by transient transfection of the pCMV-HA-I-SceI expression plasmid (Liang et al., 1998) with Jet-PEI according to the manufacturer’s instructions (Polyplus transfection), and cells were incubated for 48 hours. Cells were collected in PBS and 50 mM EDTA, pelleted and fixed with 2% paraformaldehyde for 20 minutes. The percentage of GFP-expressing cells was scored by FACS analysis using a BD Accuri C6 flow cytometer (BD Biosciences).

For silencing experiments, 20000 cells were seeded 1 day before transfection with siRNAs, using INTERFERin following the manufacturer’s instructions (Polyplus Transfection) with 20 nM of one of the following siRNAs: Control (5’-AUGAACGUGAAUUGCUCAA-3’), BRCA2-3 (5’-GCUUCAGUUGCAUAUCUUA-3’). The BRCA2 siRNA targets the 3’UTR of endogenous BRCA2 mRNA. All siRNAs were synthesized by Eurofins (France). Forty-eight hours later, the cells were transfected with the pCMV-HA-I-SceI expression plasmid. At least 3 independent experiments were performed, and HA-I-SceI expression and silencing efficiency were verified by Western blot as described below.

### Western blotting

Cells were lysed in buffer containing 20 mM Tris HCl (pH 7.5), 1 mM Na_2_EDTA, 1 mM EGTA, 150 mM NaCl, 1% (w/v) NP40, 1% sodium deoxycholate, 2.5 sodium pyrophosphate, 1 mM β-glycerophosphate, 1 mM NA_3_VO_4_ and 1 µg/ml leupeptin supplemented with complete mini protease inhibitor (Roche). Denatured proteins (20-40 µg) were electrophoresed in 9% SDS-PAGE gels or NuPAGE(tm) 3-8% Tris-Acetate Protein Gels (Invitrogen), transferred onto a nitrocellulose membrane and probed with specific antibodies: anti-BRCA2 (1/4000, ab9143, Abcam), anti-Vinculin (1/8000, ab18058 Abcam), and anti-HA (1/1,000, F-7 #sc-7392, SantaCruz). Immunoreactivity was visualized using an enhanced chemiluminescence detection kit (ECL, Pierce). The intensity of the bands was quantified by ImageJ.

### Irradiation

Cells were exposed to 0.5, 1, 2, 5, 6 or 10 Gy IR 24h after seeding using an X-ray source (1.03 Gy/min) (X-RAD 320, Precision X-Ray Inc., North Branford, CT). The cells were fixed with 4% paraformaldehyde 2h, 4h, 6h, 8h or 24h after irradiation, and immunofluorescence was performed as described below.

### Immunofluorescence

Cells were seeded onto slides, then washed with PBS, treated with CSK buffer (100 mM NaCl, 300 mM sucrose, 3 mM MgCl_2_, 10 mM Pipes pH 6.8, 1 mM EGTA, 0.2X Triton, and protease inhibitor cocktail (complete ULTRA Tablets, Roche) and fixed in 2% paraformaldehyde for 15 min. The cells were then permeabilized in 0.5% Triton-X 100 for 5 min, saturated with 2% BSA and 0.05% Tween20 and probed with anti-RAD51 antibody (1/500, PC130, Merck Millipore) for 2 h at 37°C. After 3 washes in PBS-Tween20 (0.05%) at RT, the cells were probed with Alexa-coupled anti-mouse or anti-rabbit secondary antibody (1/1,000, Invitrogen) for 1h at 37°C. After 3 washes, the cells were mounted in DAKO mounting medium containing 300 nM DAPI and visualized using a fluorescence microscope (Zeiss Axio Observer Z1) equipped with an ORCA-ER camera (Hamamatsu). Image processing and foci counting were performed using the ImageJ software.

### Statistical Analysis

Statistical analyses were performed using GraphPad Prism 3.0 (GraphPad Software).

## Acknowledgements

We thank Baptiste Fouquet for help in some experiments, Jean Soulier and the “Cellulothèque des hémopathies de l’Hôpital Saint-Louis » for the gift of FANCD1 primary fibroblasts and Alexandra Benachi for fetal ovaries. This study was supported by Université Paris Diderot (SC), Université Paris Sud-Paris Saclay (ED, AH, MM), by the Agence Nationale de Biomedecine (AH, MM) and by Institut Universitaire de France (GL). BSL was supported by the Ligue Nationale contre le cancer “Equipe labellisée 2017”, Agence Nationale de la Recherche (ANR-16-CE12-0011-02 and ANR-16-CE18-0012-02), AFM-Téléthon and Institut National du Cancer (INCa-PLBIO18-232).

## Authors contributions

MM, GL and BSL designed research studies. ED, AH, ST and SM conducted experiments. SC, ED, AH, GL, BSL and MM acquired and analysed data. MM, SC, BSL, AH and GL wrote the manuscript.

## FIGURES SUPPLEMENTS

**Figure 2–figure supplement 1.**
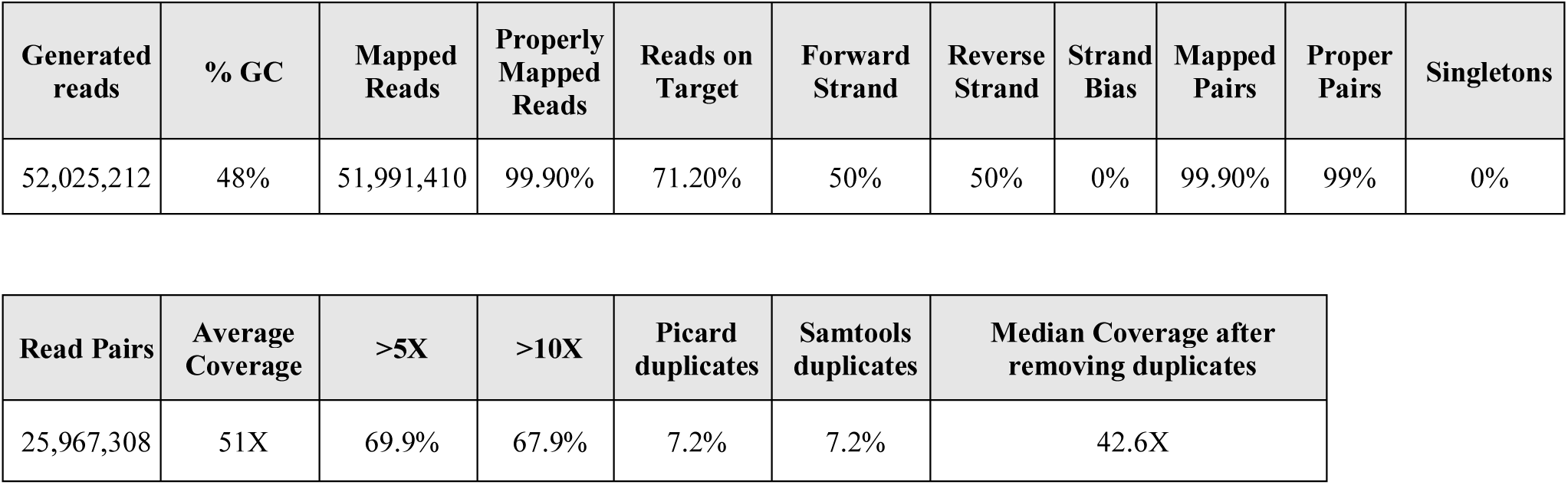
Whole Exome Sequencing and mapping data for the patient with POI.

**Figure 2–figure supplement 2.**
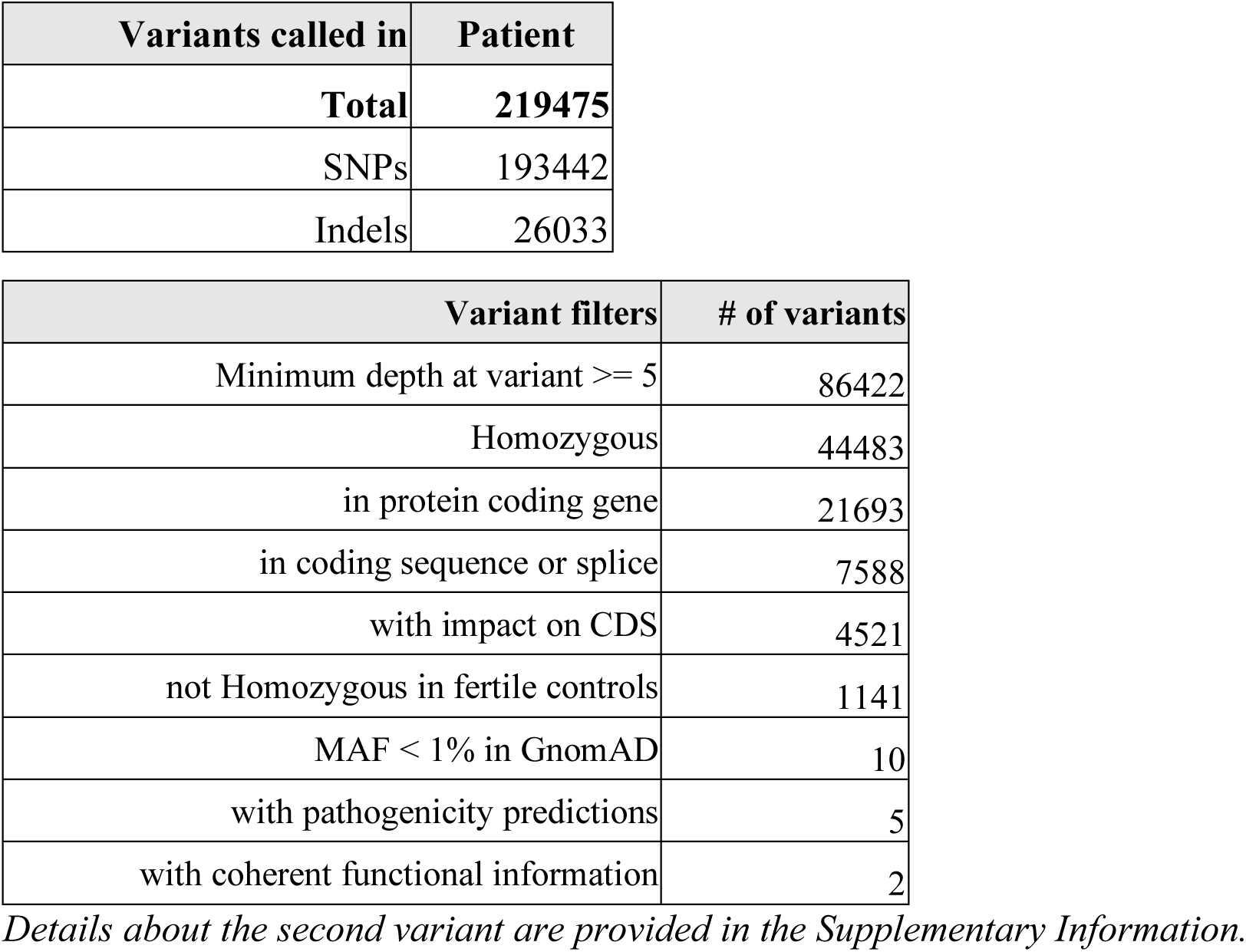
Filtering of the variants identified by Whole Exome Sequencing in the POI patient.

**Figure 2–figure supplement 3.**
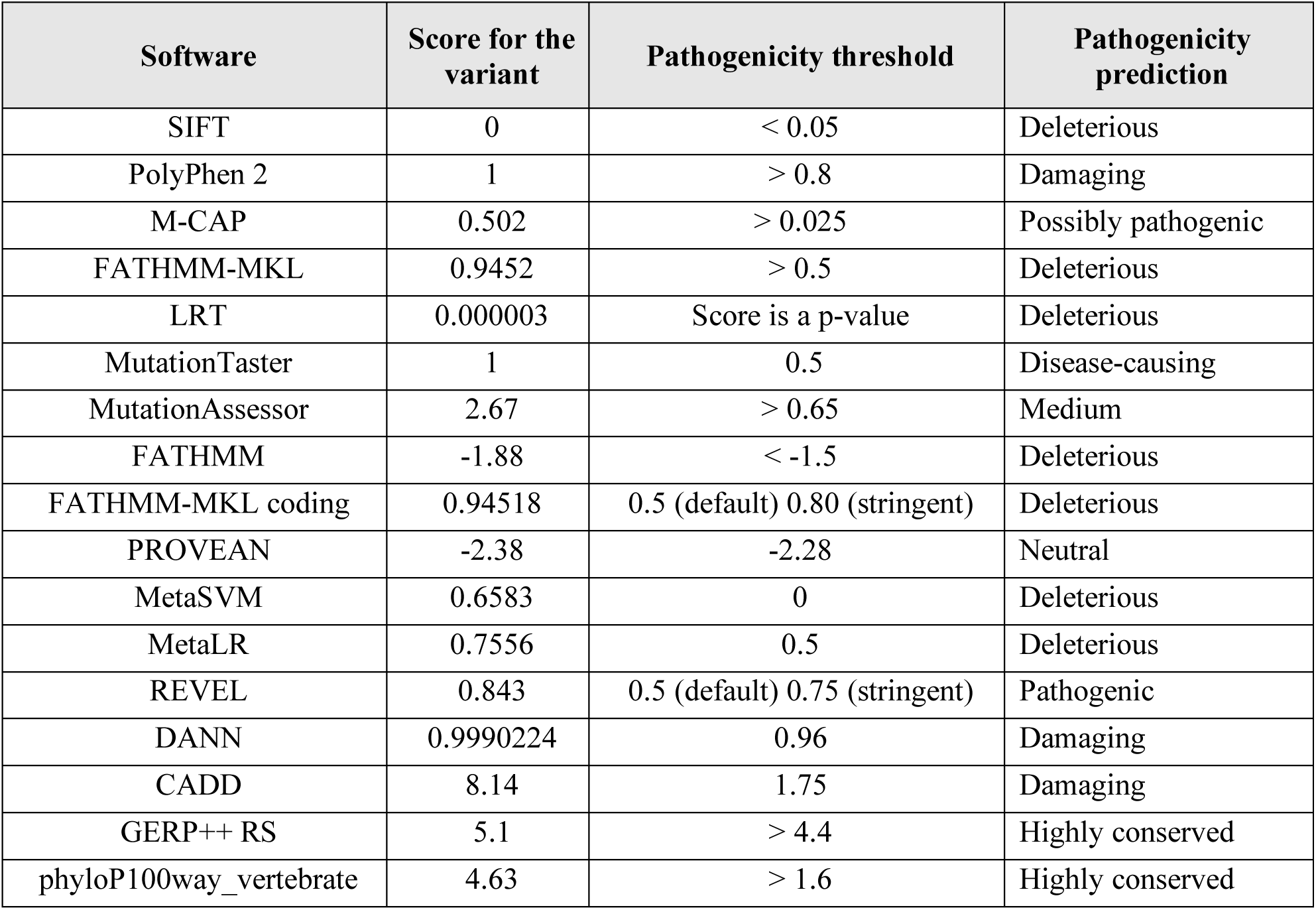
Pathogenicity predictions for the R2842C variant in *BRCA2*.

**Figure 2–figure supplement 4.**
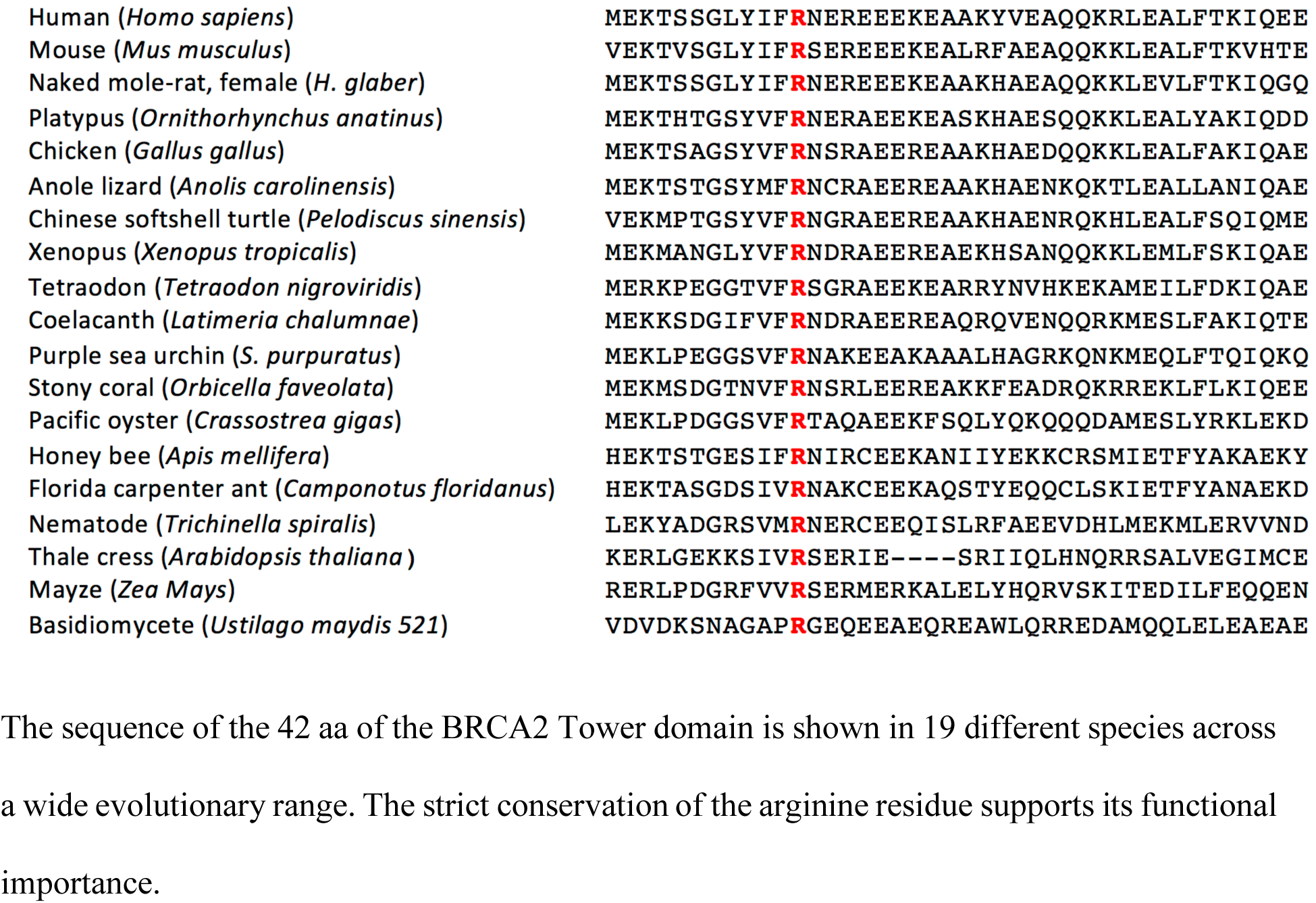
Conservation of the mutated BRCA2 Arg 2842 aminoacid across species.

**Figure 6-figure supplement 1.**
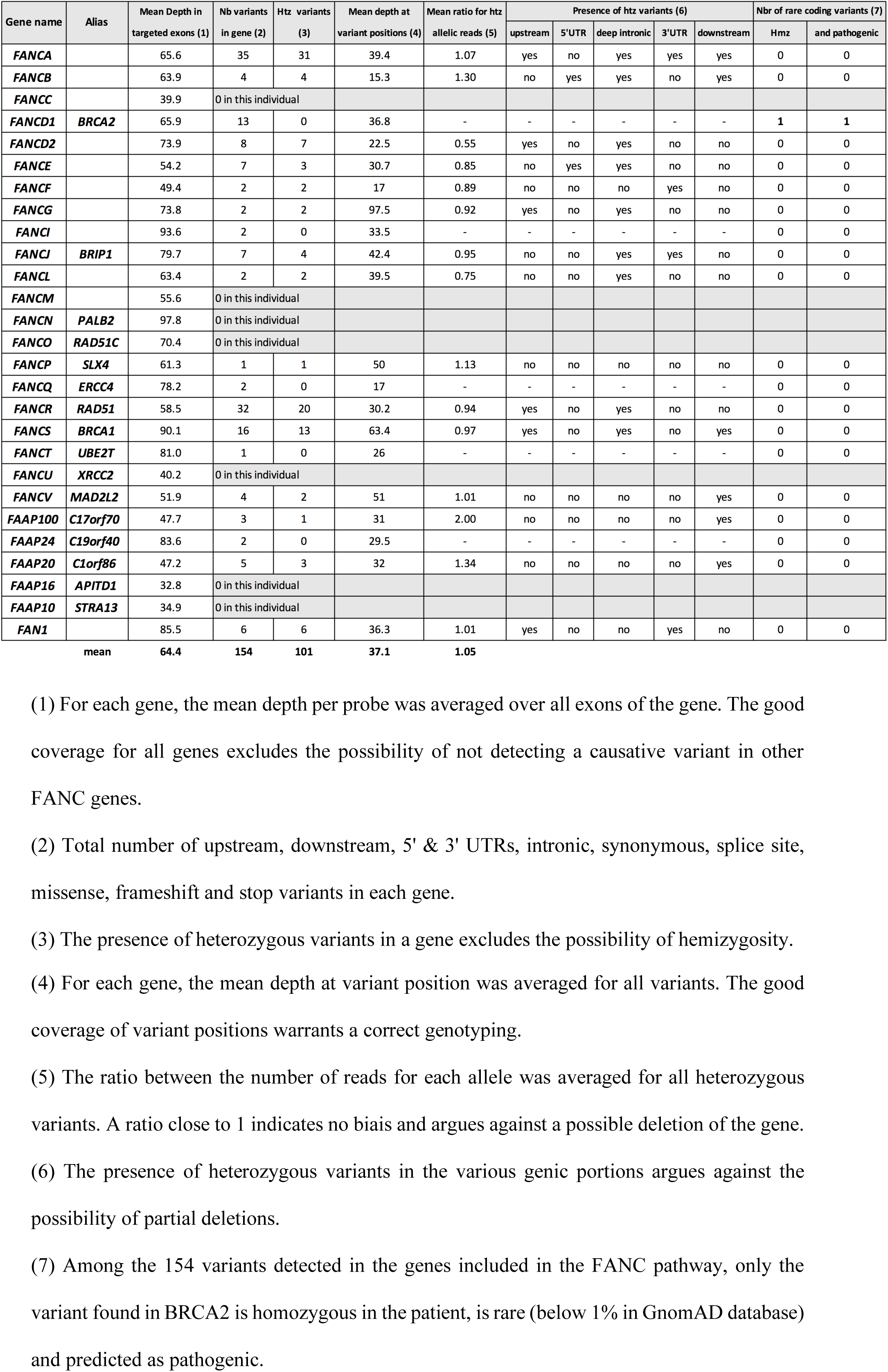
Whole Exome Sequencing data from the POI patient for all genes included in the FANC pathway, with the exception of BRCA2, to exclude a potential causative variant in all these genes.

## Supplementary Information

### Variant analysis in the patient with POI

Variants were annotated by SnpEff and VEP and were filtered on the basis of their homozygosity in the patient, their absence in unrelated fertile in-house controls and a minor allele frequency (MAF) below 0.01 in all available databases. Further filtering on available functional data for a possible role in fertility yielded only two plausible candidate variants. The first variant was a missense rs539695846 in *ARGHEF7*. This gene lies about 250 kb away from one of the 6 loci identified by genome-wide association study as influencing age at natural menopause (Stolk et al., 2009). ARHGEF7 is expressed ubiquitously with a maximum in the brain, and encodes a cytoplasmic Rho guanine nucleotide exchange factor that plays a role in cell proliferation, in particular through phosphorylation of FOXO3a (Chahdi and Sorokin, 2008). As *FOXO3a* knockout mice are infertile due to early depletion of the follicle pool (Castrillon et al., 2003), this regulation could be the basis for a possible role of *ARHGEF7* in the age of menopause. However, a recent study showed no association between the polymorphism besides *ARHGEF7* and AMH levels, a reliable marker of ovarian reserve, in childhood cancer survivors (van Dorp et al., 2013) which lessens the interest of this gene in fertility.

The second variant was the *BRCA2* missense rs80359104 characterized in this study.

### Targeted Next Generation Study in the mother

The patient’s mother had delayed conception followed by secondary amenorrhea at the age of 33 years, not investigated in Turkey. Her amenorrhea, reflecting either a central or peripherical hypogonadism, could be explained in part by obesity, known to contribute to ovulatory dysfunction and amenorrhea (Mircea et al., 2007). Thus, we performed a targeted next generation sequencing (NGS) to eliminate other genetic cause that might explain the potential precocious menopause in the mother. We obtained an average of 1 Gb of sequences with more than 98% of mappable reads and a mean depth of 150x. Nearly 96% of bases were covered to a minimum depth of 20x and more than 95% of the read bases had a Qscore of above 30. A total of four hundred and thirty-seven (437) variants were detected. Twenty variants have a frequency lower than 2% according to the ExAC base. Six false positive variants were ruled out with a careful examination of the corresponding BAMs using IGV. Of the 14 remaining variants, 8 are intronic variants with no predicted effect on splicing. Of the 6 exonic variants, one is a synonymous variant without impact on splicing. Of the remaining five exonic variants, 4 are predicted to be benign by the M-CAP prediction software (Table S1). The only remaining variant was the *BRCA2* missense mutation detected in the propositus (our patient, Figure 1, V2), *BRCA2*: c. c.8524C>T; p. Arg2842Cys.

*BRCA2* heterozygous mutations were associated with lower AMH levels, reflecting the ovarian reserve (Daum et al., 2018). We cannot exclude that the heterozygous *R2842C-BRCA2* mutation could have an impact on the mother’s ovarian reserve in addition to environmental factors.

---

**Table S1.** Rare Variants detected by Targeted Next Generation Sequencing in the mother.

| CHR | POS | REF | ALT | AF | Gene | Exon | cDNA | Protein | GnomAD<br>Genome | GnomAD<br>Exome | Kaviar | ACMG | M-CAP |
| --- | --- | --- | --- | --- | --- | --- | --- | --- | --- | --- | --- | --- | --- |
| chr13 | 32945129 | C | T | 0.500 | BRCA2<br>(NM_000059.3) | exon20 | c.8524C>T | p.Arg2842Cys | 3.24e-05 | 8.14e-06 | . | Pathogenic<br>Supporting | Possibly<br>pathogenic |
| chr5 | 140071240 | C | G | 0.500 | HARS2<br>(NM_012208.3) | exon1 | c.7C>G | p.Leu3Val | 0.00352 | 0.00291 | 0.000643 | VUS | Likely benign |
| chr12 | 53818988 | T | G | 0.500 | AMHR2<br>(NM_020547.2) | exon4 | c.464T>G | p.Phe155Cys | . | . | . | VUS | Likely benign |
| chr22 | 31867903 | C | T | 0.500 | EIF4ENIF1<br>(NM_019843.3) | exon3 | c.97G>A | p.Glu33Lys | 0.00452 | 0.00378 | 0.0038 | Pathogenic<br>Supporting | Likely benign |
| chr8 | 31015010 | A | G | 0.500 | WRN<br>(NM_000553.4) | exon33 | c.3946A>G | p.Ile1316Val | . | 0.000134 | 0.000161 | VUS | Likely benign |

\* VUS: Variant of unknown significance

